## Supplementary information for "Structure of hepatitis B/D antiviral drug Bulevirtide bound to its receptor protein NTCP"

**This PDF file includes:**
Supplementary Tables 1-2
Supplementary Figures 1-7
Supplementary References

**Supplementary Table 1. Cryo-EM data collection and processing.**

| **Sample** | NTCP-BLV- Fab3- and Nb |
|---|---|
| PDB | 8RQF |
| EMDB | EMD-19440 |
| **Data collection and processing** | |
| Microscope, | Titan Krios G3i, |
| Electron gun, | X-FEG, |
| Energy filter, | GIF BioQuantum Gatan K3 |
| Detector, | |
| Voltage (kV) | 300 |
| Magnification | 130,000x |
| Electron exposure per movie ($e^-/Å^2$) | 55 |
| Defocus range (μm) | -0.5 to -2.5 |
| Pixel size (Å) | 0.648 |
| Numbers of micrographs | 2,610 |
| Symmetry imposed | C1 |
| Initial particle images (no.) | 1,696,766 |
| Final particle images (no.) | 128,700 |
| Map resolution (Å) at FSC threshold of 0.143 | 3.41 |
| Map sharpening *B* factor ($Å^2$) | 118.0 |
| Processing software | CryoSPARC v4.4.0 |
| **Refinement** | |
| Initial model used (PDB code) | 7ZYI |
| FSC threshold (0.143/0.5) | 3.4/3.6 |
| Model composition | |
| Non-hydrogen atoms | 6,950 |
| Protein residues | 903 |
| Ligands | LIG: 1 |
| B factor ($Å^2$) | |
| Protein | 22.96/177.66/83.01 |
| Ligand | 44.34/48.12/48.12 |
| R. m. s. d deviations | |
| Bond lengths (Å) | 0.003 |
| Bond angles (°) | 0.640 |
| **Validation** | |
| MolProbity score | 1.81 |
| Clash score | 9.09 |
| Poor rotamers (%) | 0.26 |
| Ramachandran plot | |
| Favored (%) | 95.30 |
| Allowed (%) | 4.70 |
| Disallowed (%) | 0.00 |

**Supplementary Table 2. List of interactions between BLV and NTCP**

| BLV peptide | | NTCP |
|---|---|---|
| No. | AA | |
| 2 | Myr-G | F128, C129, L131, G132, M133, D152, Y156, K157 |
| 3 | T | D152, K153, V272 |
| 4 | N | D152, K153, V154, P155, Y156, T268, I269, V272 |
| 5 | L | K157, G158 |
| 6 | S | P155, G158, I159, Q264, T268 |
| 7 | V | G158, I161, S162 |
| 8 | P | P154, G158, I159, S162, N262, Q264, L265 |
| 9 | N | L31, V32, L35, N262, Q264 |
| 10 | P | L35, G102, N103, S162, V166, N262, V263, M290 |
| 11 | L | L31, M34, L35, I38, L104 |
| 12 | G | L31, V263, Q264, M290 |
| 13 | F | L27, L31, V202, S206, V263, Q264, S267, F283, P286, L287, M290 |
| 14 | F | F18, D24, L27, V210, Q264, S267, F283, F284, L287 |
| 15 | P | Q264, S267, T268, N271, F283 |
| 16 | D | G19, D24, S28, Q264 |
| 17 | H | D24, L27, S28, L31, Q264 |
| 18 | Q | V32, Q264, T268 |
| 19 | L | V32, L35, S162, L165, V166 |
| 20 | D | - |
| 21 | P | - |
| 22 | A | - |
| 23 | F | L25, S28, V29, V32 |
| 24 | G | - |
| 25 | A | G19 |
| 26 | N | G19, K20, D24, L25, S28 |
| 27 | S | G19, K20 |
| 28 | N | K20 |
| 29 | N | - |
| 30 | P | K20 |
| 31 | D | - |
| 32 | W | T268, N271, V272 |
| 33 | D | N271, V272 |
| 34 | F | - |
| 35 | N | K153, V272, A273 |
| 36 | P | - |
| 37 | N | - |
| 38 | K | N271, V272, A273, F274, P275, P276 |
| 39 | D | Y146, A273, F274, P275, V278 |
| 40 | H | P275, E277, V278 |

| | | |
|---|---|---|
| 41 | W | N87, I88, L91, I145, Y146, D147, F274, V278 |
| 42 | P | - |
| 43 | E | N87, I145, D147 |
| 44 | A | K86, N87, I88, V278 |
| 45 | N | K86 |
| 46 | K | L85, K86, N87 |
| 47 | V | R84, L85, K86, N87 |
| 48 | G | N87, G144, I145 |

### Supplementary Figures

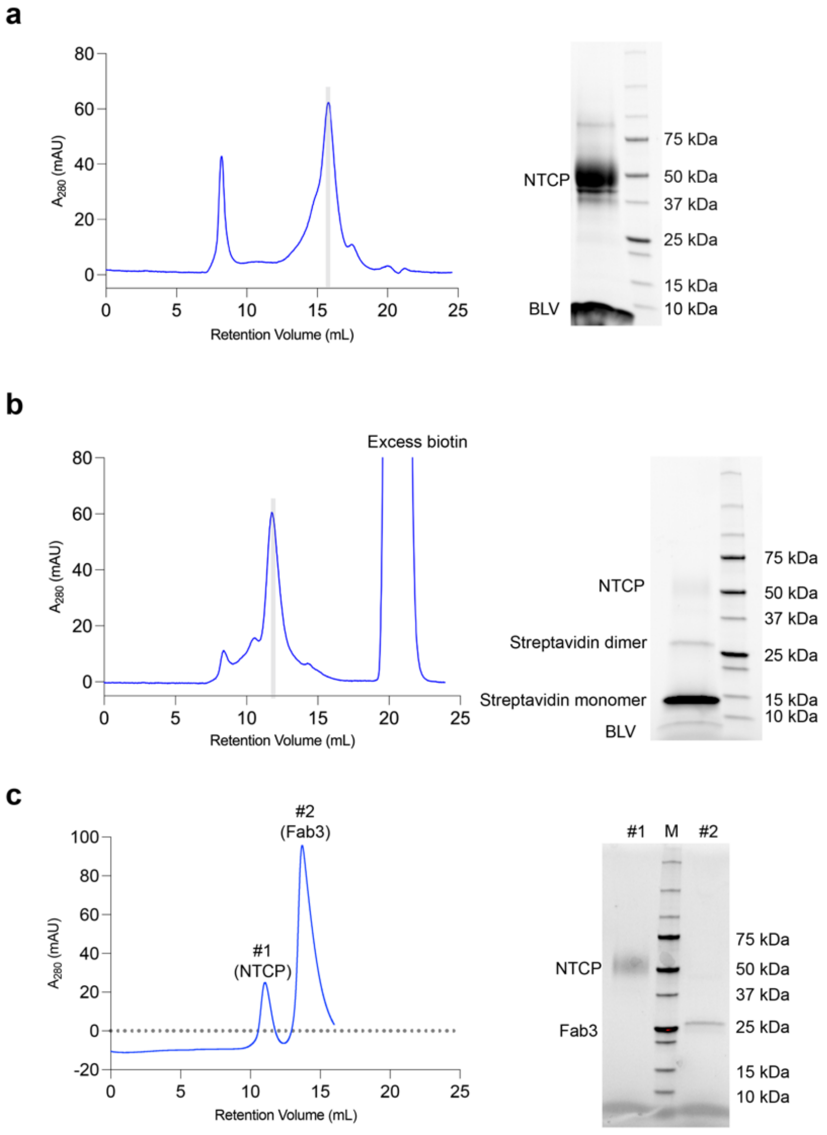

**Supplementary Fig. 1. Purification of NTCP-BLV complex. a** Left: Size-exclusion chromatogram (SEC) of detergent-solubilized NTCP bound to BLV. Right: SDS-PAGE analysis of highlighted fraction. **b** Left: SEC of NTCP-BLV complex after biotinylation. Highlighted fraction was used for pull-down assay. Right: SDS-PAGE analysis of a sample immobilized on streptavidin-coated paramagnetic particles (see Method section). **c** Left: SEC of NTCP and Fab3 mixture in the absence of BLV, confirming lack of complex formation. Right: SDS-PAGE analysis of selected fractions

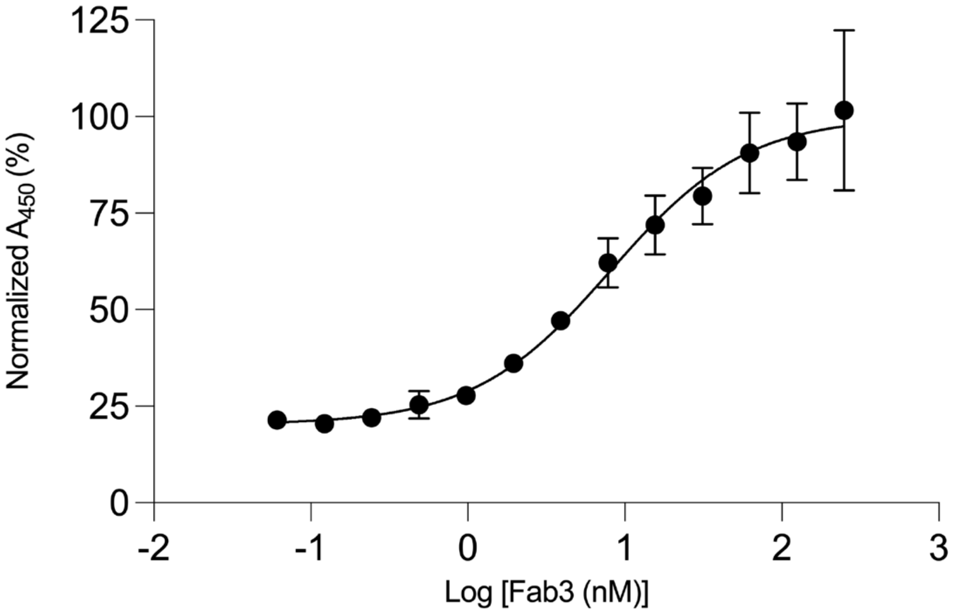

**Supplementary Fig. 2. Estimation of the binding affinity ($EC_{50}$) of Fab3 to BLV-bound NTCP in detergent.** Normalized protein ELISA at varying concentrations of Fab3 to determine the binding affinity, $EC_{50}$, of Fab3 to detergent-solubilized BLV-bound NTCP. The $EC_{50}$ was calculated to be 8 nM. Data points represent the mean, error bars indicate the standard deviation of three independent measurements.

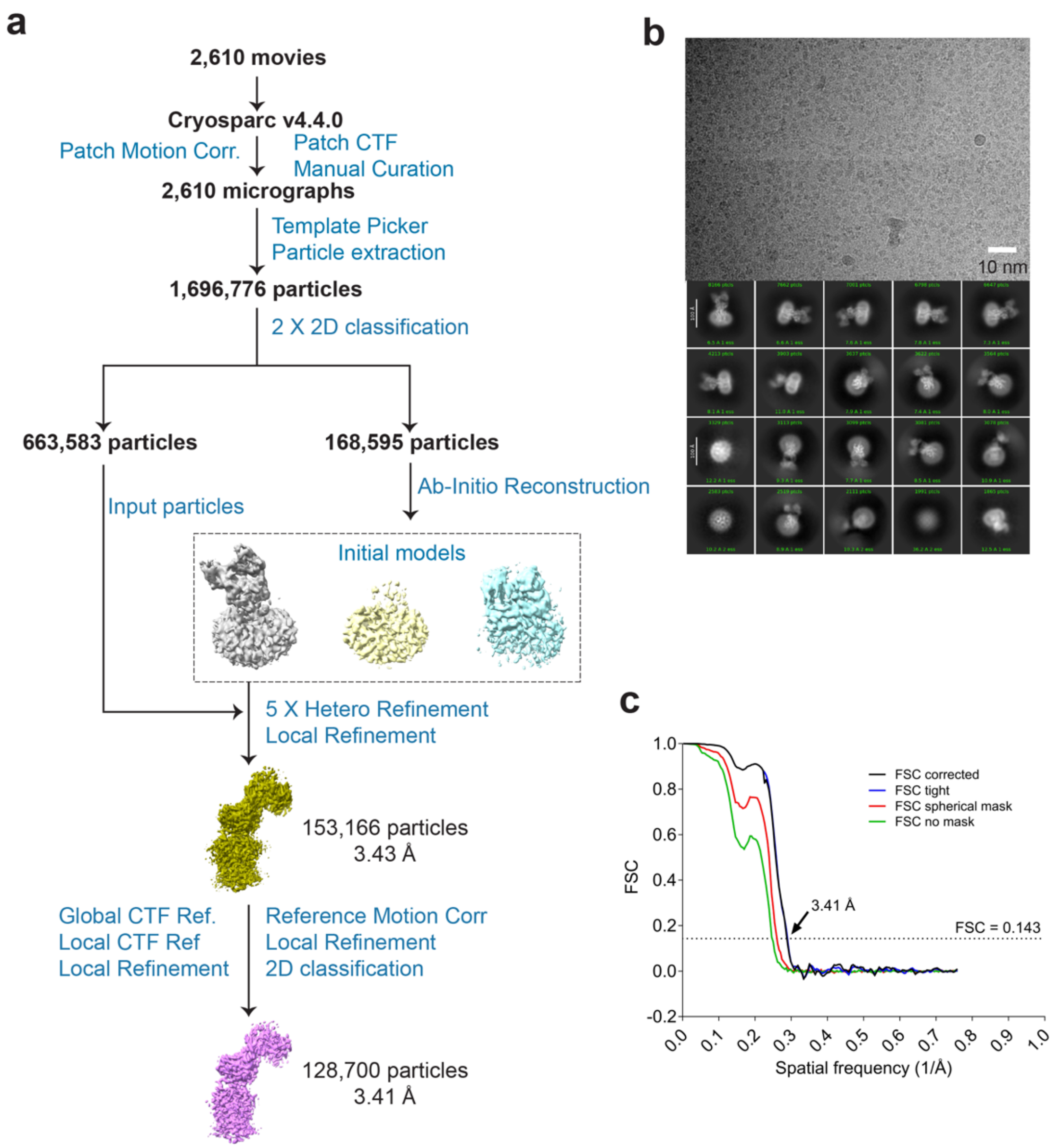

**Supplementary Fig 3. Cryo-EM data processing of the NTCP-BLV-Fab3-Nb complex. a** Data processing flowchart **b** Representative motion-corrected micrograph and 2D class averages of BLV-bound NTCP-Fab3-Nb complex. **c** Fourier shell correlation (FSC) curves.

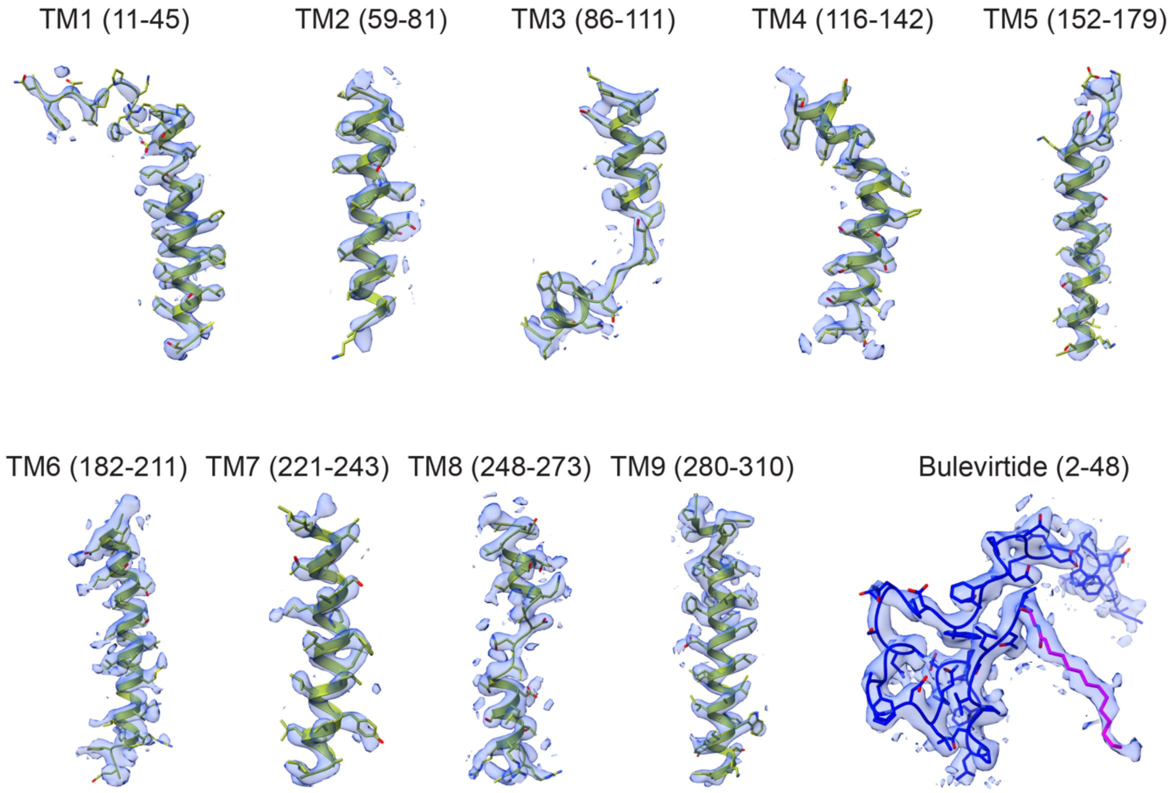

**Supplementary Fig. 4. Cryo-EM density of NTCP-BLV-Fab3-Nb complex.** The TM helices of NTCP (green) and the Bulevirtide peptide (blue) are shown as ribbons and numbered. The corresponding EM density is displayed as a blue transparent surface.

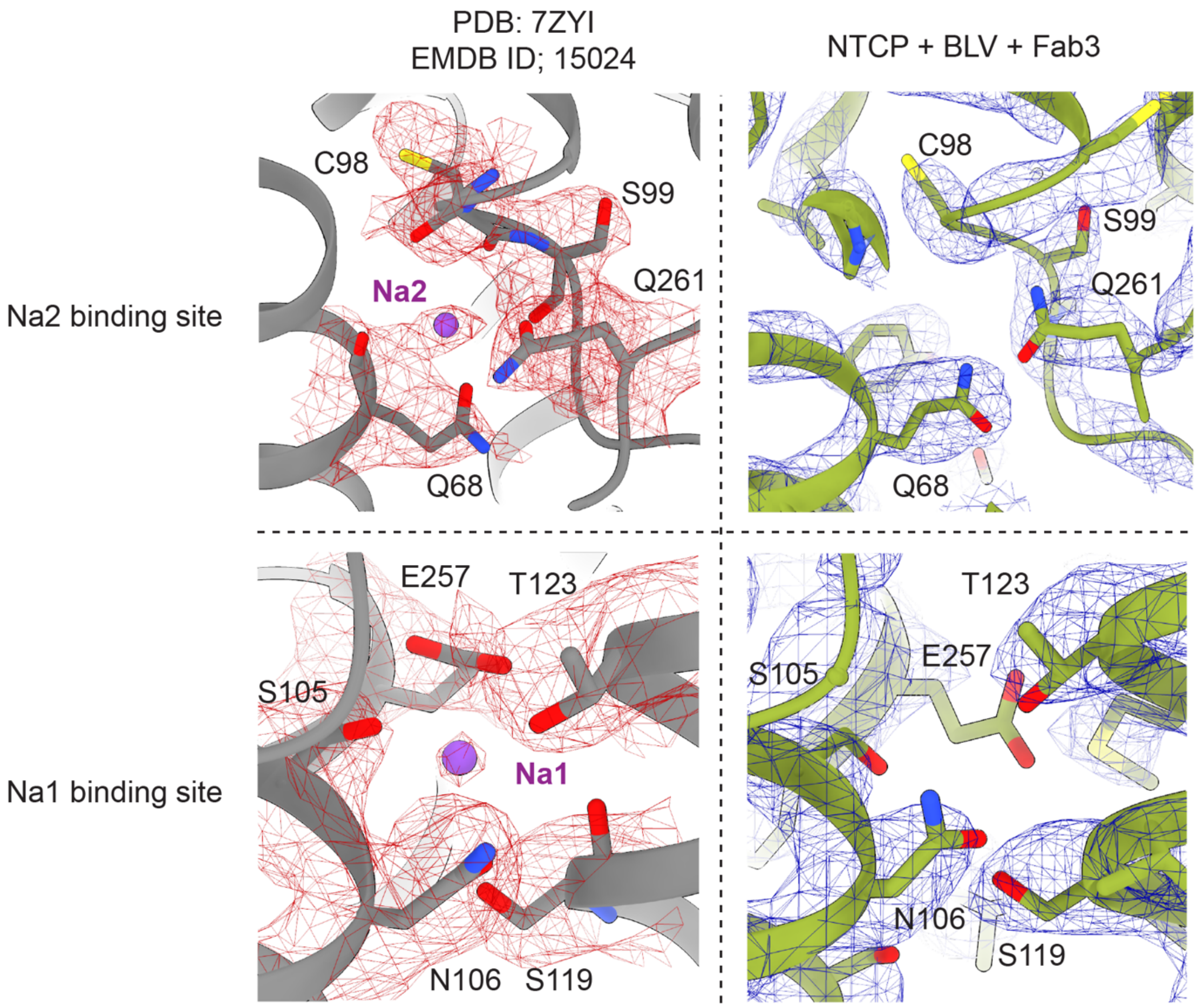

**Supplementary Fig. 5. Comparison of sodium binding sites in BLV-bound and substrate-bound NTCP.** The structure of substrate-bound NTCP (PDB ID: 7ZYI, EMDB ID: 15024) is shown in grey ribbon representation, and residues coordinating bound sodiums (Na1 and Na2) are shown in stick representation and labeled. The EM density is shown as red mesh. The structure of BLV-bound NTCP (this study) is shown in green ribbon representation, residues are labeled and the EM density is shown as blue mesh.

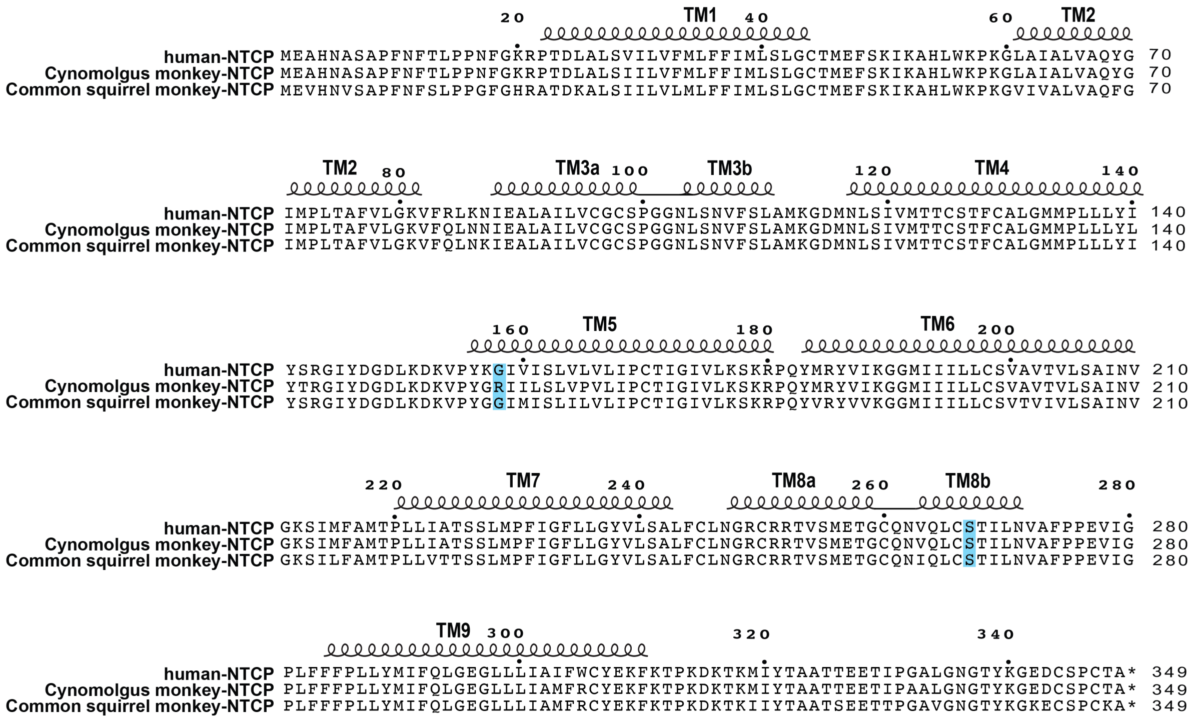

**Supplementary Fig. 6. NTCP conservation among primates.** Amino acid sequence alignment of NTCP from human, cynomolgus monkey (representative of Old World monkeys), and common squirrel monkey (representative of New World monkeys). Residues highlighted in blue are discussed in the text.

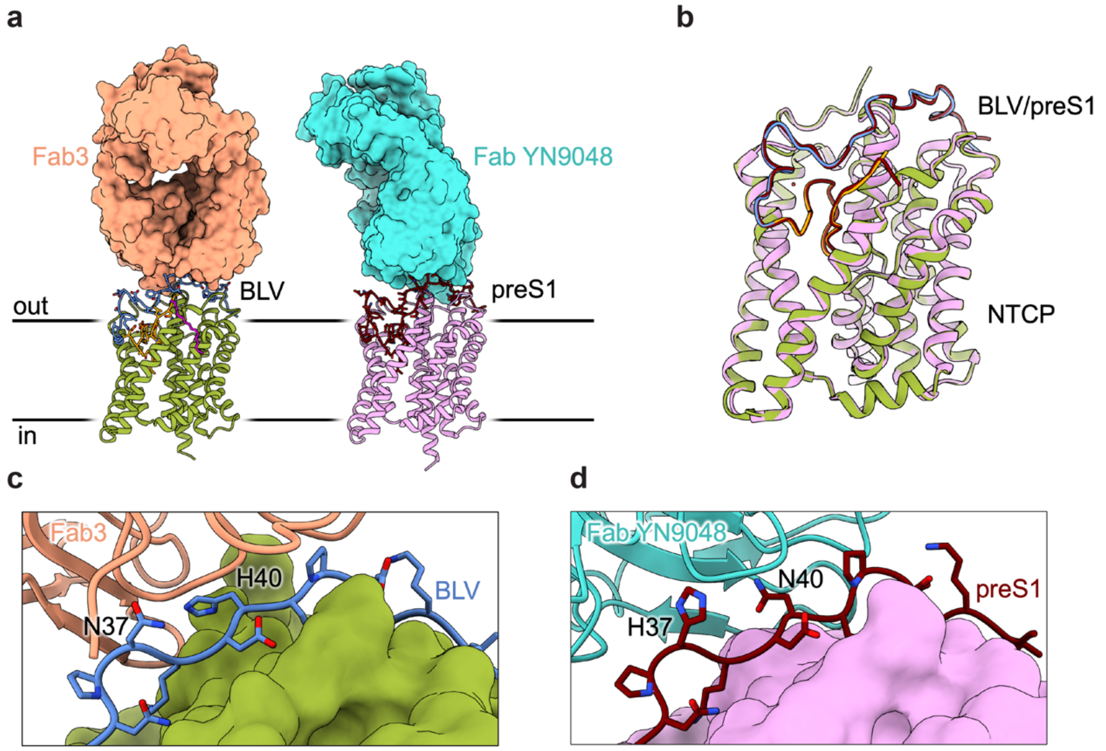

**Supplementary Fig. 7. Comparison of BLV-bound and preS1-bound NTCP structures. a** Structure of NTCP-BLV-Fab3-Nb (this study) and NTCP-preS1-Fab YN9048 (PDB ID: 8HRX)[1]. NTCP is displayed in ribbon representation, Fabs as surface, and bound peptides as sticks. **b** Superposition of BLV-bound and preS1-bound NTCP structures, colored as in panel **a**. **c-d** Close-up view of the peptides interacting with the corresponding Fabs. Two residues involved in the binding to each Fab differ between BLV (**c**) and preS1 (**d**), these are labeled.
